# Stage-specific gene and transcript dynamics in human male germ cells

**DOI:** 10.1101/2022.03.22.485121

**Authors:** Lara M. Siebert-Kuss, Henrike Krenz, Tobias Tekath, Marius Wöste, Sara Di Persio, Nicole Terwort, Margot J. Wyrwoll, Jann-Frederik Cremers, Joachim Wistuba, Martin Dugas, Sabine Kliesch, Stefan Schlatt, Frank Tüttelmann, Jörg Gromoll, Nina Neuhaus, Sandra Laurentino

## Abstract

Cell differentiation processes are highly dependent on cell stage-specific gene expression, including timely production of alternatively spliced transcripts. One of the most transcriptionally rich tissues is the testis, where the process of spermatogenesis, or generation of male gametes, takes place. To date, germ cell-specific transcriptome dynamics remain understudied due to limited transcript information emerging from short-read sequencing technologies. To fully characterize the transcriptional profiles of human male germ cells and to understand how the human spermatogenic transcriptome is regulated, we compared whole transcriptomes of men with different types of germ cells missing from their testis. Specifically, we compared the transcriptomes of testis lacking germ cells (Sertoli cell-only phenotype; SCO; n=3), with an arrest at the stage of spermatogonia (SPG; n=4), spermatocytes (SPC; n=3), and round spermatids (SPD; n=3), with the transcriptomes of testis with normal and complete spermatogenesis (Normal; n=3). We found between 839 and 4,138 differentially expressed genes (DEGs, log_2_ fold change ≥ 1) per group comparison, with the most prevalent changes observed between SPG and SPC arrest samples, corresponding to the entry into meiosis. We detected highly germ cell-type specific marker genes among the topmost DEGs of each group comparison. Moreover, applying state-of-the-art bioinformatic analysis we were able to evaluate differential transcript usage (DTU) during human spermatogenesis and observed between 1,062 and 2,153 genes with alternatively spliced transcripts per group comparison. Intriguingly, DEGs and DTU genes showed minimal overlap (< 8%), suggesting that stage-specific splicing is an additional layer of gene regulation in the germline. By generating the most complete human testicular germ cell transcriptome to date, we unravel extensive dynamics in gene expression and alternative splicing during human spermatogenesis.

## Introduction

Human male germ cell differentiation is a complex process requiring cell type-specific transcriptome regulation. Disturbances in spermatogenesis causing male infertility range from maturation arrest at different germ cell stages to complete lack of germ cells, known as Sertoli cell-only (SCO) phenotype. Although an increasing number of male infertility cases can be attributed to pathogenic variants in genes involved in spermatogenesis (Houston *et al*., 2021), the number of causative pathogenic variants identified so far remains small (Tüttelmann *et al*., 2018). The identification and understanding of genetic causes for male infertility is hindered by the lack of data regarding transcriptomic dynamics during human spermatogenesis.

In order to obtain testicular cell-type specific gene expression profiles, previous studies took advantage of samples with distinct histological phenotypes of male infertility using samples matched by cellular composition (Winge *et al*., 2018) or by performing comparative microarray analyses of samples differing in the presence of one specific germ cell-type (von Kopylow *et al*., 2010; Chalmel *et al*., 2012; Lecluze *et al*., 2018). For example, comparing testicular tissues with SCO and spermatogonial arrest phenotypes, which only differ in the presence of spermatogonia, von Kopylow *et al*., (2010) were able to identify transcripts specifically expressed by spermatogonia. The authors identified the spermatogonial markers *FGFR3* and *UTF1*, which are currently considered specific markers for different spermatogonial subpopulations (Guo *et al*., 2018; Sohni *et al*., 2019; Di Persio *et al*., 2021). Chalmel *et al*. (2012) expanded on this approach by including samples from different developmental stages and arrest phenotypes, thereby extracting the transcriptional profiles of additional germ cell types. These studies demonstrated that the comparison of distinct arrest phenotypes allows the identification of transcripts expressed at specific stages of germ cell differentiation during normal spermatogenesis (von Kopylow *et al*., 2010; Chalmel *et al*., 2012). Technological developments such as RNA sequencing (RNA-seq) now enable an unbiased and more comprehensive analysis of the transcriptome. Specifically, single cell RNA-sequencing (scRNA-seq) of human testicular tissues has revolutionized germ cell-specific RNA profiling by allowing the identification of cell type-specific gene expression patterns (Guo *et al*., 2018; Hermann *et al*., 2018; Wang *et al*., 2018; Sohni *et al*., 2019; Di Persio *et al*., 2021). However, scRNA-seq results in sparser data compared to conventional bulk RNA-seq and, by sequencing from the poly-A tail of transcripts, generates limited information on transcriptional isoforms (Tekath and Dugas, 2021). Total RNA-seq therefore results in the most complete capture of the transcriptome, including all transcripts obtained through post-transcriptional processing. The testis presents unusual high levels of these post-transcriptional events, including alternative splicing (AS) (Kan *et al*., 2005). AS enables the production of different transcripts and proteins from a single gene, thereby also constituting a crucial regulatory mechanism for gene expression. For example, storage of immature mRNAs allows protein synthesis at transcriptionally silent stages of mouse spermatogenesis (Iguchi *et al*., 2006; Naro *et al*., 2017). During human male germ cell differentiation, AS events have so far been understudied, with the exception of the association between hormone receptor genes splice site variants and human male infertility (Song *et al*., 2002; Bruysters *et al*., 2008). Knowledge of the changes in isoforms that result from AS during human spermatogenesis would open a new avenue for identifying so far unknown causes for male infertility.

In this study, we aimed at generating the most complete human testicular germ cell transcriptome to date. Combining the advantages of scRNA-seq data and total RNA-seq of distinct pathological phenotypes, and using sophisticated bioinformatic analyses, we unveiled the transcriptional profiles of male germ cell types and determined the changes in AS patterns during human male germ cell differentiation.

## Materials and Methods

### Ethical approval

Male infertility patients included in this study underwent surgery for microdissection testicular sperm extraction (mTESE; n=15) or to rule out a suspected malignant tumor (n=1) at the Department of Clinical and Surgical Andrology of the Centre of Reproductive Medicine and Andrology, University Hospital of Münster, Germany. Each patient gave written informed consent (ethical approval was obtained from the Ethics Committee of the Medical Faculty of Münster and the State Medical Board no. 2008-090-f-S) and one additional testicular sample for the purpose of this study was obtained. Tissue proportions were snap-frozen or fixed in Bouin’s solution.

### Patient selection

In this study, we included testicular biopsies with a homogenous histological phenotype in both testes from men showing SCO (SCO-1/ M1045, SCO-2/ M911, SCO-3/M1742), spermatogenic arrests at the spermatogonial (SPG-1/ M1570, SPG-2/ M1575, SPG-3/ M1072, SPG-4/ M2822), spermatocyte (SPC-1/ M1369, SPC-2/ M799, SPC-3/ M921), and round spermatid stage (SPD-1/ M2227, SPD-2/ M1311, SPD-3/ M1400) (Table I). We excluded patients with germ cell neoplasia and a history of cryptorchidism as well as acute infections. For complete representation of the spermatogenic process, samples with qualitatively and quantitatively normal spermatogenesis were included in this study (Normal-1/M1544, Normal-2/M2224, Normal-3/M2234) obtained from patients with obstructive azoospermia, e.g. due to congenital bilateral absence of the *vas deferens* (CBAVD; Normal-1), anorgasmia (Normal-2) or due to suspected tumor that was not confirmed (Normal-3). Prior to surgery, all patients underwent physical evaluation, hormonal analysis of luteinizing hormone (LH), follicle stimulating hormone (FSH), and testosterone (T), and semen analysis (World Health Organization, 2010). In addition to conventional karyotyping and screening for azoospermia factor (AZF) deletions, whole exome sequencing (WES) was performed for all patients, except for SPG-4 (who had undergone chemotherapy because of leukemia) and one with normal spermatogenesis (Normal-3). WES data were generated within the Male Reproductive Genomics (MERGE) study as previously published (Wyrwoll *et al*., 2020) and were screened for variants in 230 candidate genes that have at least a limited level of evidence for being associated with male infertility according to a recent review (Houston *et al*., 2021). We also included a screening in the recently published genes *ADAD2, GCNA, MAJIN, MSH4*, MSH5, *RAD21L1, RNF212, SHOC1, STAG3, SYCP2, TERB1, TERB2*, and *TRIM71*, which are associated with non-obstructive azoospermia (Riera-Escamilla *et al*., 2019; Krausz *et al*., 2020; Schilit *et al*., 2020; Hardy *et al*., 2021; Salas-Huetos *et al*., 2021; Torres-Fernández *et al*., 2021; Wyrwoll *et al*., 2021). We screened for rare (minor allele frequency [MAF] in gnomAD database < 0.01), possibly pathogenic variants (stop-, frameshift-, and splice site variants) with a read depth > 10x, that were detected in accordance with the reported mode of inheritance.

**Table 1:** Clinical characteristics of the patient groups. Data are presented as mean ± standard deviation. Percentage of tubules with elongated spermatids (%ES), round spermatids (%RS), spermatocytes (%SPC), spermatogonia (%SPG), Sertoli cell-only phenotype (%SCO), and tubular shadows (%TS). Score refers to Bergmann and Kliesch score (Bergmann and Kliesch, 2010). Hormonal parameters for follicle stimulating hormone (FSH), luteinizing hormone (LH) and testosterone (T). ^a^Patient SPD-3 had a low number of XXY karyotype mosaicism (47,XXY[2]/46,XY[28]). ^b^TESE results: N-1 had 100/100 sperm, N-2 had an average of 89/100 sperm; No TESE result available for N-3 due to consultation to exclude a malignant tumor. SCO – Sertoli cell-only; SPG – spermatogonial arrest; SPC – spermatocyte arrest; SPD – spermatid arrest; Normal – normal spermatogenesis; n.d. – not determined.

| Patient groups | Karyotype | Histological parameters of tubules |  |  |  |  |  |  | Hormonal parameters (normal range) |  |  | Sperm mTESE |
| --- | --- | --- | --- | --- | --- | --- | --- | --- | --- | --- | --- | --- |
|  |  | Score | % ES | % RS | % SC | % SG | % SCO | % TS | FSH (1-7U/l) | LH (2-10U/l) | T(>12nmol/l) |  |
| SCO (n=3) | 46,XY | 0 | 0 | 0 | 0 | 0 | 98.7 (± 1.5) | 1.3 (± 1.5) | 13.3 (± 4.2) | 5.8 (± 2.6) | 13.7 (± 3.4) | No |
| SPG (n=4) | SPG-1, SPG-2,<br>SPG-3: 46,XY;<br>SPG-4: n.d. | 0 | 0 | 0 | 0 | 31.0 (±34.6) | 34.3 (± 20.7) | 35.0 (± 20.6) | 20.4 (± 14.2) | 13.4 (± 9.7) | 16.2 (± 6.9) | No |
| SPC (n=3) | 46,XY | 0 | 0 | 0 | 89.3 (± 11.0) | 4.7 (±4.6) | 1.0 (± 1.0) | 5.3 (± 5.5) | 5.7 (± 1.3) | 5.7 (± 4.5) | 9.9 (± 2.4) | No |
| SPD (n=3) | SPD-1, SPD-2:<br>46,XY; SPD-3:<sup>a</sup> | 0 | 0 | 28.3 (± 2.3) | 59.3 (± 18.0) | 3.0 (±2.0) | 1.7 (± 2.9) | 8.7 (± 14.2) | 7.4 (± 0.9) | 3.7 (± 0.5) | 18.7 (± 5.7) | No |
| Normal (n=3) | 46,XY | 8-10 | 87.3 (± 8.6) | 3.3 (± 2.5) | 8.7 (± 5.7) | 0 | 0 | 1.0 (± 1.0) | 2.5 (± 1.3) | 2.6 (± 1.0) | 24.7 (± 2.2) | Yes<sup>b</sup> |

### Histological evaluation of the human testicular biopsies

After overnight fixation in Bouin’s solution, the tissues were washed in 70% ethanol, embedded in paraffin, and sectioned at 5 μm. AppiClear (Applichem, Cat# A4632.2500) was used to dewax the tissue section. The cellular composition of all testicular biopsies (n=16) was histologically examined on two periodic acid-Schiff (PAS)-stained sections from two independent biopsies per testis. For PAS staining, the sections were first incubated with 1% PA (Sigma-Aldrich, Cat# 1.005.240.100) and then in Schiffs reagent (Sigma-Aldrich, Cat# 1.090.330.500). Cell nuclei were counterstained with Mayer’s hematoxylin solution (Sigma-Aldrich, Cat# 1.092.490.500). After washing in tap water and dehydration through increasing ethanol concentrations and AppiClear, slides were closed with Merckoglas (Sigma-Aldrich, Cat# 1.039730.001). The slides were scanned using the Precipoint Viewpoint software (Precipoint, Freising, Germany). The biopsies were evaluated based on the Bergmann and Kliesch scoring method (Bergmann and Kliesch, 2010), which assigns a score from 0 to 10 to each patient according to the percentage of tubules containing elongated spermatids. Furthermore, the percentage of the seminiferous tubules with round spermatids, spermatocytes or spermatogonia as the most advanced germ cell type was assessed, as well as seminiferous tubules with SCO or hyalinized tubules (tubular shadows) (Table I).

### RNA extraction from testicular tissues

We extracted total RNA from snap-frozen testicular tissues from all biopsies using the Direct-zol™ RNA Microprep kit (Zymo Research, CA, USA) according to manufacturer’s protocol. Quantity and quality of isolated RNA were evaluated using RNA ScreenTape and the TapeStation Analysis software 3.1.1 (Agilent Technologies, Inc., CA, USA). All samples had intact ribosomal 18S and 21S bands. Samples with an RNA integrity number (RIN) >3.6 were included in the analysis (Suntsova *et al*., 2019).

### Library preparation and sequencing

Next-generation sequencing was performed by the service unit Core Facility Genomics of the medical faculty at the University of Münster. Libraries were prepared according to the NEBNext Ultra RNA II directional Library Prep kit (New England Biolabs, MA, USA) after NEBNext rRNA depletion (New England Biolabs, MA, USA). The NextSeq HO Kit (Illumina Inc., CA, USA) with 150 cycles was used for paired end sequencing on the NextSeq 500 system (Illumina Inc., CA, USA) with ~400 Million single reads per run.

### Data processing

We processed the raw sequence data with the nextflow analysis pipeline nf-core/rnaseq 2.0 (Ewels *et al*., 2020) and annotated the transcripts with GENCODE release 36 genome annotation based on the GRCh38.p13 genome reference (Frankish *et al*., 2019). Gene expression counts were estimated using *Salmon* (Patro *et al*., 2017).

### Differential gene expression analysis

All data were analyzed within the R Statistical Environment (RCoreTeam, 2020). We used DESeq2 (Love *et al*., 2014) for analyzing differentially expressed genes (DEGs) following the standard workflow for *Salmon* quantification files. DESeq2 uses a generalized linear model based on estimated size factors and dispersion to calculate the log_2_ fold changes for each gene (Love *et al*., 2014). Annotation was performed using the biomaRt R package Normalization was performed using DESeq2 with the median of ratios method (Love *et al*., 2014). Genes with a total count > 10 were considered for further analysis. DEGs were calculated for each group comparison, i.e. SCO vs. SPG, SPG vs. SPC, SPC vs. SPD, and SPD vs. Normal. *P*-values are calculated based on Wald test and adjusted with Benjamini-Hochberg. Genes with a false discovery rate (FDR) < 0.05 and a log_2_ fold change (FC) ≥ 1 were considered DEGs. Dispersion of samples was visualized using DESeq2’s *PCAplot* function for the top 500 genes with a total count > 10.

To evaluate gene expression of selected genes of interest at single-cell level, we generated uniform manifold approximation and projection (UMAP) plots (McInnes *et al*., 2020) based on our previously published dataset (Di Persio *et al*., 2021) using the tool Seurat (Stuart *et al*., 2019; Hao *et al*., 2021).

### Differential transcript usage analysis

For computing differential transcript usage (DTU) we employed the R package DTUrtle (Tekath and Dugas, 2021), following the vignette workflow for human bulk RNA-seq analysis. As for the DEG analysis, we annotated the transcripts with GENCODE release 36 genome annotation. We calculated DTU genes for each group comparison (i.e. SCO vs. SPG, SPG vs. SPC, SPC vs. SPD, and SPD vs. Normal) with the *run_drimseq* function. DTUrtle conducts statistical analyses based on DRIMSeq (Nowicka and Robinson, 2016), i.e. a likelihood ratio test is used on the estimated transcript proportions and precision parameter (Tekath and Dugas, 2021). To increase the statistical power of the analysis, we filtered out transcripts with low impact, i.e. less than 5% usage for all samples or a corresponding total gene expression of less than 5 counts for all samples before the statistical testing. Also, only genes with at least two high impact transcripts were considered. From the analysis we obtained genes with an overall significant change in transcript usage as well as the corresponding transcripts that drive the change in usage in those genes (both with overall FDR < 0.05).

To decrease the number of analyzed transcripts per DTU genes, a post-hoc filtering was applied, i.e. transcripts whose proportional expression deviated by less than 10% between samples were excluded. In this study, we decided to only include transcripts, which fulfill the criterion that all samples of one group must have a higher transcript usage compared to all samples of the other group.

### Pathway Analysis

Molecular function of DEGs and DTU genes were assessed via the Ingenuity Pathway Analysis software (IPA; Qiagen, Hilden, Germany). A Benjamini-Hochberg multiple testing correction *P*-value (FDR) <0.01 was used as threshold for significant molecular functions in IPA. We selected the top 20 significant terms for molecular functions.

### Statistical analysis

Statistical analysis was conducted as described in sections for differential gene expression analysis, differential transcript usage analysis, and pathway analysis.

## Results

### Clinical characteristics of the study cohort

Hormonal evaluation revealed that patients with normal spermatogenesis had FSH values within the reference range, whereas most patients with spermatogenic arrests had elevated FSH levels (Table I). Other than patient SPD-3, who had a low grade XXY mosaicism (47,XXY[2]/46,XY[28]), no patients showed chromosomal abnormalities. By analyzing WES data of our patients with unknown reasons for infertility, we did not identify any likely high impact pathogenic variants in known male infertility candidate genes.

### Testicular phenotypes are recapitulated at RNA level

To obtain whole transcriptome expression profiles, we sequenced total RNA of human testicular biopsies with SCO, SPG, SPD, as well as normal spermatogenesis (n=16) (Fig. 1A). Prior to sequencing, a careful histological examination (Fig. 1B) ensured that both testes presented comparable phenotypes, and no sperm was found via mTESE, except in the normal samples (Table I). Following total RNA-seq, principal component analysis (PCA) organized the spermatogenic arrest samples in consecutive order (Fig. 1C), mirroring their sequential spermatogenic phenotypes.

**Figure 1:**
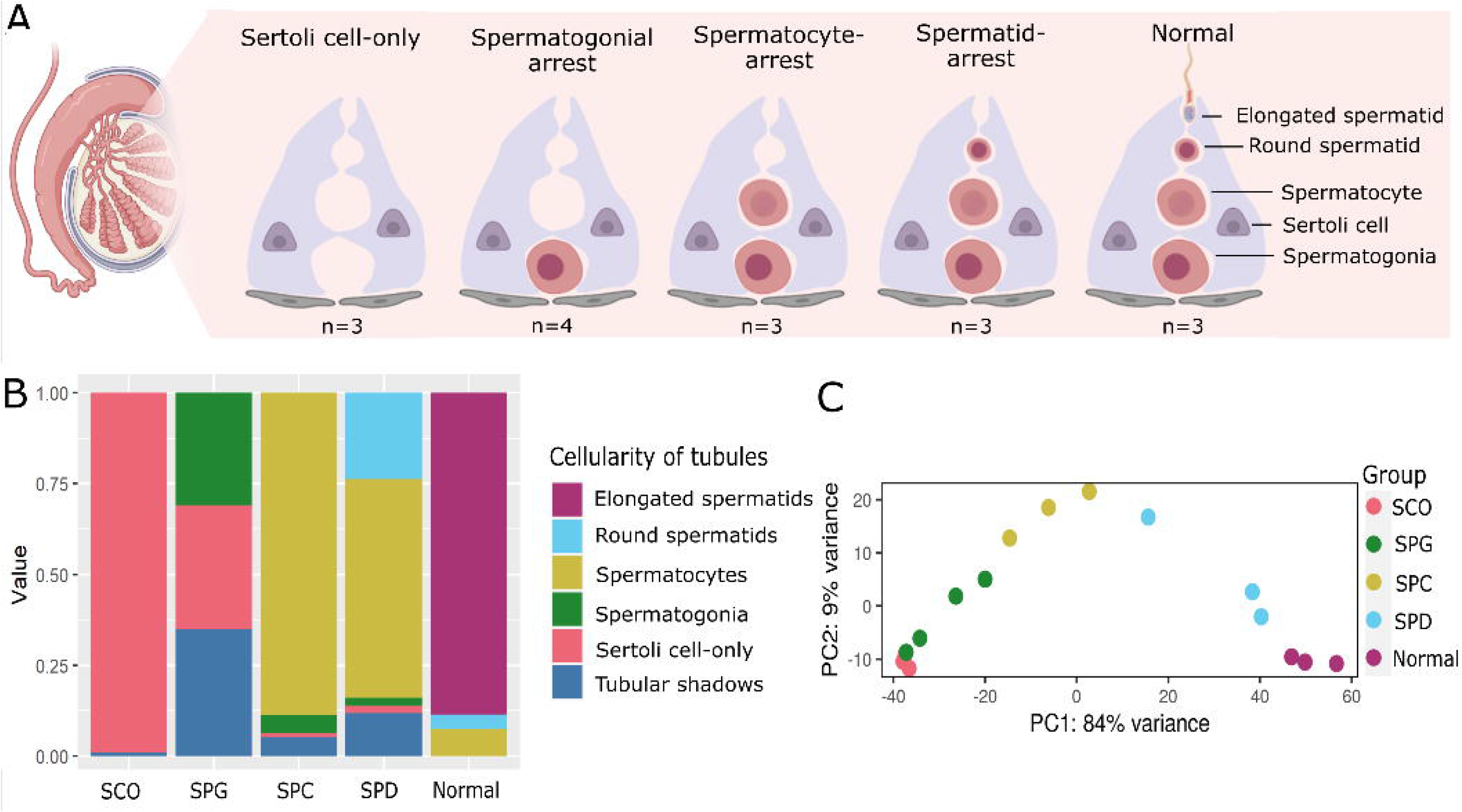
Cellular composition of the human testicular biopsies. (A) Schematic illustration depicts the cellular composition of the testicular biopsies with Sertoli cell-only (SCO) arrest at the spermatogonial (SPG), spermatocyte (SPC) and spermatid (SPD) stage as well as samples with normal spermatogenesis (Normal). (B) Stacked barplots represent the proportion of round seminiferous tubules and their most advanced germ cell-type in each sample group. The cellularity of samples from one group is averaged. (C) A principal component analysis (PCA) plot depicts clustering of the total RNA sequenced samples based on the top 500 genes.

### Comparative analysis reveals germ cell-specific transcriptional profiles

We aimed at generating germ cell-specific expression profiles to study transcriptional changes throughout spermatogenesis. To this end we performed differential gene expression analysis between groups of different cellularity: SCO versus SPG (comparison 1), SPG versus SPC (comparison 2), SPC versus SPD (comparison 3) and SPD versus normal (comparison 4). This revealed between 839 and 4,138 DEGs in the four comparisons calculated (FDR < 0.05 and absolute log_2_ FC ≥ 1, Fig. 2). In the SCO versus SPG comparison, most transcriptional changes were due to the increased expression of 2,073 genes in SPG samples (Fig. 2A, Supplementary Table SI). Co-expression of DEGs among all samples revealed the level of gene expression remained high in the other groups containing spermatogonia (SPC, SPD, Normal), indicating that most of these transcripts originate from the presence of spermatogonia. Indeed, among the highly expressed genes were well-known spermatogonial genes such as *MAGEA4* and *FGFR3* (Supplementary Table SII). The most prominent changes in gene expression were found when comparing SPG with SPC samples (Fig. 2B, Supplementary Table SIII). The 2,886 genes that were high in expression included spermatocyte-specific genes like *AURKA* and *OVOL1* (Supplementary Table SII). The same genes also showed high expression in SPD and normal samples and low to absent expression in SPG and SCO. This indicates that these genes are specific to spermatocytes, rather than the result of gene expression alterations in other cell types. When comparing SPC with SPD samples we found 2,345 highly expressed genes in SPD samples (Fig. 2C, Supplementary Table SIV), including spermiogenesis marker genes *TNP1* and *PRM1* (Supplementary Table SII). These genes also showed higher expression in normal samples and lower expression in samples lacking spermatids (SPC, SPG, SCO), in accordance with their spermatid-specific expression pattern. The most subtle changes in gene expression were detected when comparing SPD with samples showing normal spermatogenesis (Supplementary table V), in which the presence of elongated spermatids is the only histological difference. Genes with increased expression in normal samples (776) showed lower expression levels in the spermatogenic arrest samples (SPD, SPC, SPG, SCO) (Fig. 2D) and, among others, included genes associated with the sperm flagellum like *CATSPER3* and *TEKT2* (Supplementary Table SII).

**Figure 2:**
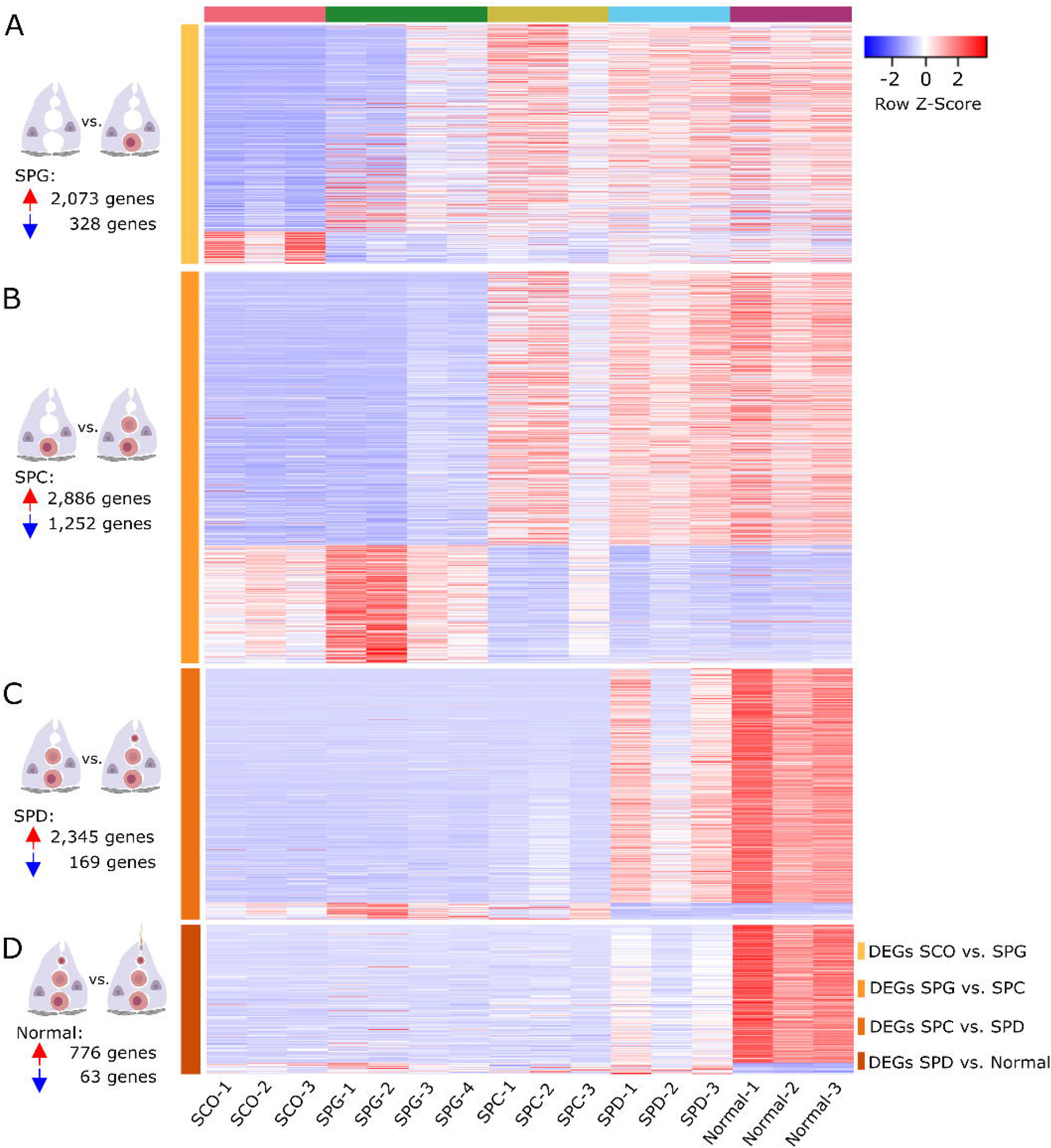
Co-expression of the DEGs among all samples. (A-D) Heatmaps display the normalized expression counts of the DEGs (rows) of the (A) SCO vs. SPG, (B) SPG vs. SPC, (C) SPC vs. SPD, and (D) SPD vs. Normal group comparisons across all samples (columns) scaled via a row Z-score. Red = increased; blue = decreased.

### Novel germ cell-specific marker genes and their expression at single cell resolution

To identify novel germ cell-specific marker genes, we focused on the top 100 DEGs per group comparison with elevated expression in SPG, SPC, SPD, and normal samples. After evaluating the expression of all genes for their germ cell-specificity in our published scRNA-seq dataset of 3 patients with normal spermatogenesis (Di Persio *et al*., 2021) (Fig. 3A), we show 3 genes per group comparison as examples. Accordingly, from the SCO vs SPG comparison, we selected the *leucine zipper protein 4* gene (*LUZP4), testis specific protein Y-linked 4 (TSPY4*), and *anomalous homeobox (ANHX*), which showed increased expression in SPG samples (Fig. 3B). Importantly, at single cell level, the expression of these genes was specific for spermatogonia (Fig. 3C). Based on the SPG vs SPC comparison we selected the *proline rich acidic protein 1 (PRAP1), ferritin heavy chain like 17 (FTHL17*) and *synaptogyrin 4 (SYNGR4*) (Fig. 3D). The spermatocyte-specific expression of these genes was confirmed in the single cell dataset (Fig. 3E). For SPD samples, genes with high expression were *proline rich 30 (PRR30), actin like 7A (ACTL7A*), and *high mobility group box 4 (HMGB4*) (Fig. 3F). Based on the expression patterns at single cell level, *PRR30, ACTL7A* and *HMGB4* were expressed in early and late spermatids (Fig. 3G). *TP53 target 5 (TP53TG5), 3-oxoacid CoA-transferase 2 (OXCT2*), and *hemogen (HEMGN*) were the highest expressed genes in normal samples in comparison to SPD samples (Fig. 3H), and also at single-cell level their expression was specific for late spermatids (Fig. 3I).

**Figure 3:**
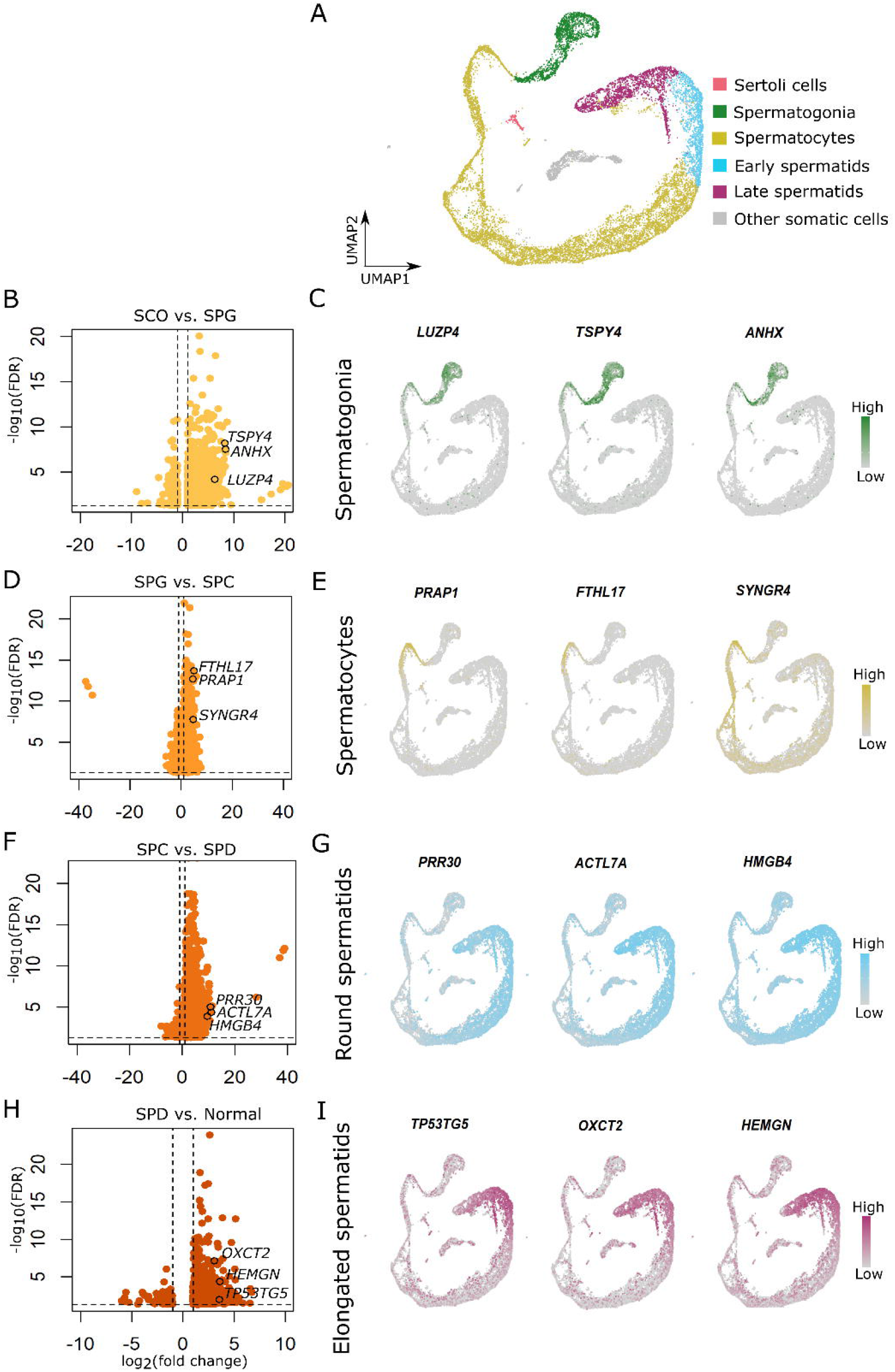
Examination of germ cell-type specific gene expression at single cell level. (A) UMAP plot depicts 15,546 cells integrated from three patients with obstructive azoospermia and normal spermatogenesis. Sertoli cell, spermatogonia, spermatocyte, early and late spermatid clusters are color-coded, respectively. (B, D, F, H) Vulcano plots of the increased and decreased genes in samples with (B) spermatogonial arrest, (D) spermatocyte, (F) and spermatid arrest, as well as in (H) normal spermatogenesis. (C, E, G, I) Feature plots show the expression of three genes selected for (C) spermatogonia, (E) spermatocytes, (G) round spermatids, (I) and elongated spermatids at single-cell level.

### Alternative splicing is uncoupled from gene expression

To study alternative splicing, we performed a DTU analysis between all four group comparisons. DTU analysis calculates and compares the proportional contributions (referred to as ‘usage’) of transcripts to the overall expression of a gene. A gene has a DTU event, i.e. is a DTU gene, when at least two of its transcripts are differentially used between two groups. We found between 1,062 and 2,153 DTU genes in each of the four comparisons (Supplementary Tables SVI-SIX). By comparing DTU genes to DEGs, we found an overlap of less than 8% in all four comparisons, indicating that the expression of most genes is regulated either at the pre- or the post-transcriptional level (Fig. 4), and that only few genes are regulated at these two levels. Furthermore, we found that the proportion of DEGs to DTUs in all group comparisons was 2:1 (Fig. 4A-C), except for SPD vs Normal, where this ratio was inversed with more DTU genes than DEGs (Fig. 4D).

**Figure 4:**
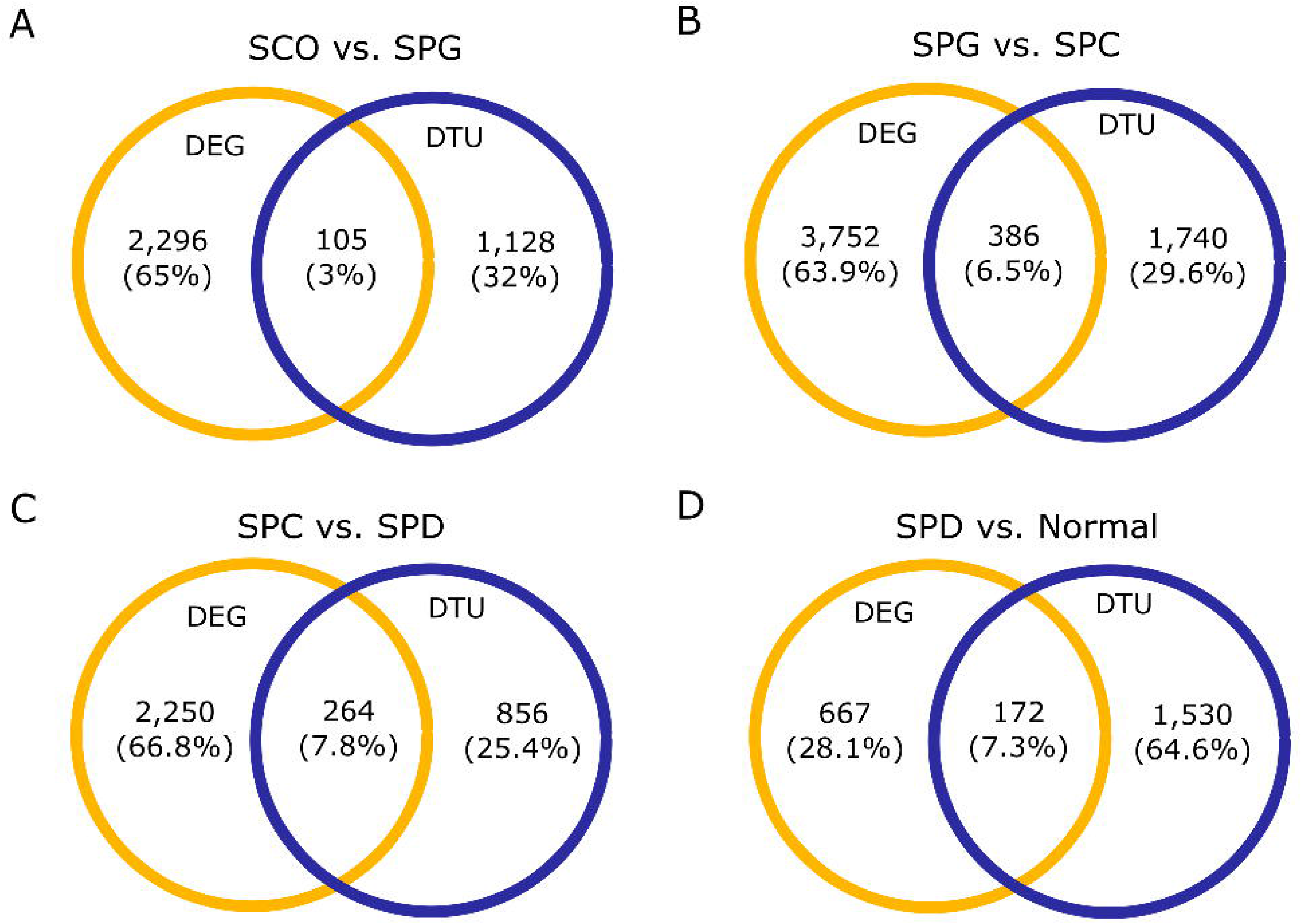
Comparison of DEG and DTU gene numbers in all four group comparisons. (A-D) Venn-diagrams display number and proportion of genes that are differentially expressed, have a DTU event, or both in the (A) SCO vs. SPG, (B) SPG vs. SPC, (C) SPC vs. SPD, and (D) SPD vs. Normal group comparisons. Yellow = DEGs, blue = DTU genes.

### DEGs and DTU genes are involved in different biological pathways

We used IPA to evaluate the molecular functions of the DEGs and DTU genes at the different germ cell stages. In line with the small overlap between the DEG and DTU gene sets, we found minor overlaps between the top 20 significantly enriched molecular functions of DEGs and DTU genes in all four groups (Fig. 5). Both gene sets contained genes involved in organization of cytoskeleton/cytoplasma, microtubule dynamics, apoptosis, necrosis, and segregation of chromosomes. IPA analysis on DEGs highlighted functional enrichment annotations that can be attributed to the most advanced germ cell type in each group comparison (e.g. development of stem cells, segregation of chromosomes) (Fig. 5A).

**Figure 5:**
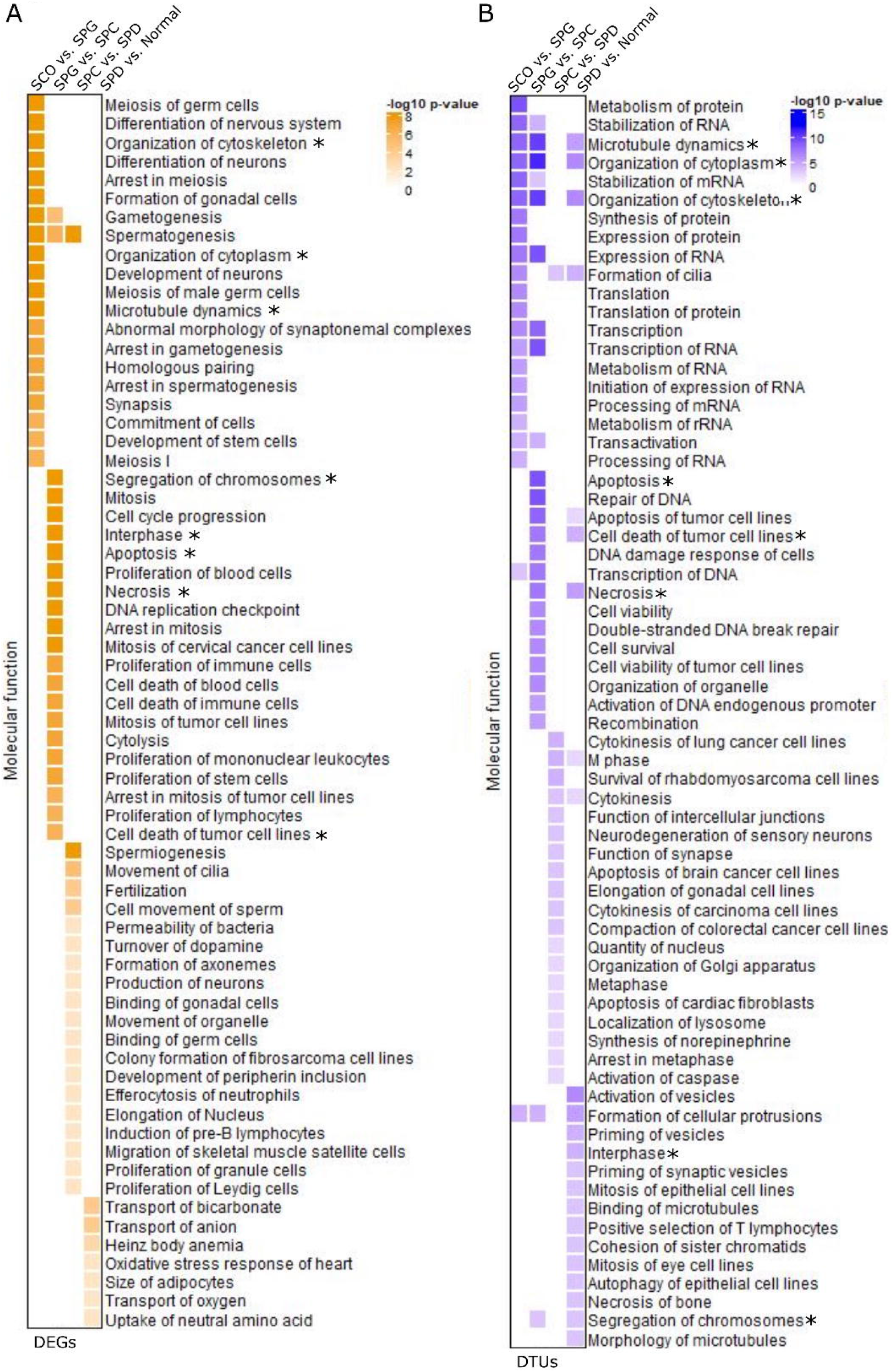
Molecular functions of DEG and DTU genes. Heatmaps of color-coded −log_10_ *p*-values display the molecular functions of (A) DEGs and (B) DTU genes per group comparison. Top 20 molecular functions with *p*-values <0.01 are included. (*) Molecular functions enriched in both, the DEG and DTU gene sets.

In comparison to the functional annotations of DEGs, 26% of molecular functions of the DTU genes overlapped across the four group comparisons (Fig. 5B). Among the overlapping terms were microtubule dynamics, organization of cytoplasm, and cytoskeleton. More general biological functions (e.g. RNA metabolism, cell survival) were enriched among the DTU genes in each group comparison.

### Stage-specific splicing is an additional layer of gene regulation in the germline

To study alternatively spliced transcripts, we investigated the transcript biotypes of selected DTU genes. In comparison to the proportional distribution of transcript biotypes annotated in GENCODE (Frankish *et al*., 2019) we found that most of the DTU events, regardless of the group comparison, result in protein coding transcripts (Fig. 6A). In the comparison between SPD arrest and normal, two protein-coding isoforms of *actin like 6A (ACTL6A*) displayed differential usage (Fig. 6B). While *ACTL6A-202* (ENST00000429709.7) was the predominant isoform, with an average usage of 52% in SPD samples, normal samples predominantly used the *ACTL6A-203* isoform (ENST00000450518.6), which has an alternative 5’ splice site (Fig. 6B). In comparison, *spermatogenesis associated 4 (SPATA4*) also showed a switch in usage for its protein coding isoforms *SPATA4-201* (ENST00000280191.7) and *SPATA4-203* (ENST00000515234.1) in the comparison of SPC versus SPD samples (Fig. 6C). SPC samples showed a significantly decreased usage of *SPATA4-201* and a significantly increased usage of *SPATA4-203*, whereas SPD samples exclusively used the *SPATA4-201* isoform (Fig. 6C). These two isoforms use alternative transcriptional start and stop sites. In contrast to *ACTL6A*, *SPATA4* was also a DEG in this group comparison and had a higher expression level in SPD samples (Supplementary Fig. S1A and S1B). Intriguingly, the second largest group of biotypes with DTU events were retained introns (Fig. 6A). For *synaptonemal complex protein 3* (*SYCP3*), we found a significantly increased usage of the retained intron isoform *SYCP3-204* (ENST00000478139.1) in SPG samples, whilst SPC samples had an increased usage of the protein coding isoform *SYCP3-202* (ENST00000392924.2) (Fig. 6D). In this group comparison, *SYCP3* showed increased expression in SPC samples (Fig. S1C). A switch in usage from coding to non-coding transcripts was also observed for *marker of proliferation Ki-67 (MKI67*) (Fig. 6E), which did not show changes in gene expression (Supplementary Fig. S1D). However, the protein coding isoform *MKI67-202* (ENST00000368654.8) was lower expressed in SPC samples in comparison to SDP samples. In contrast, the processed transcript isoform *MKI67-205* (ENST00000484853.1) showed significantly increased usage in SPC samples and decreased usage in SDP samples.

**Figure 6:**
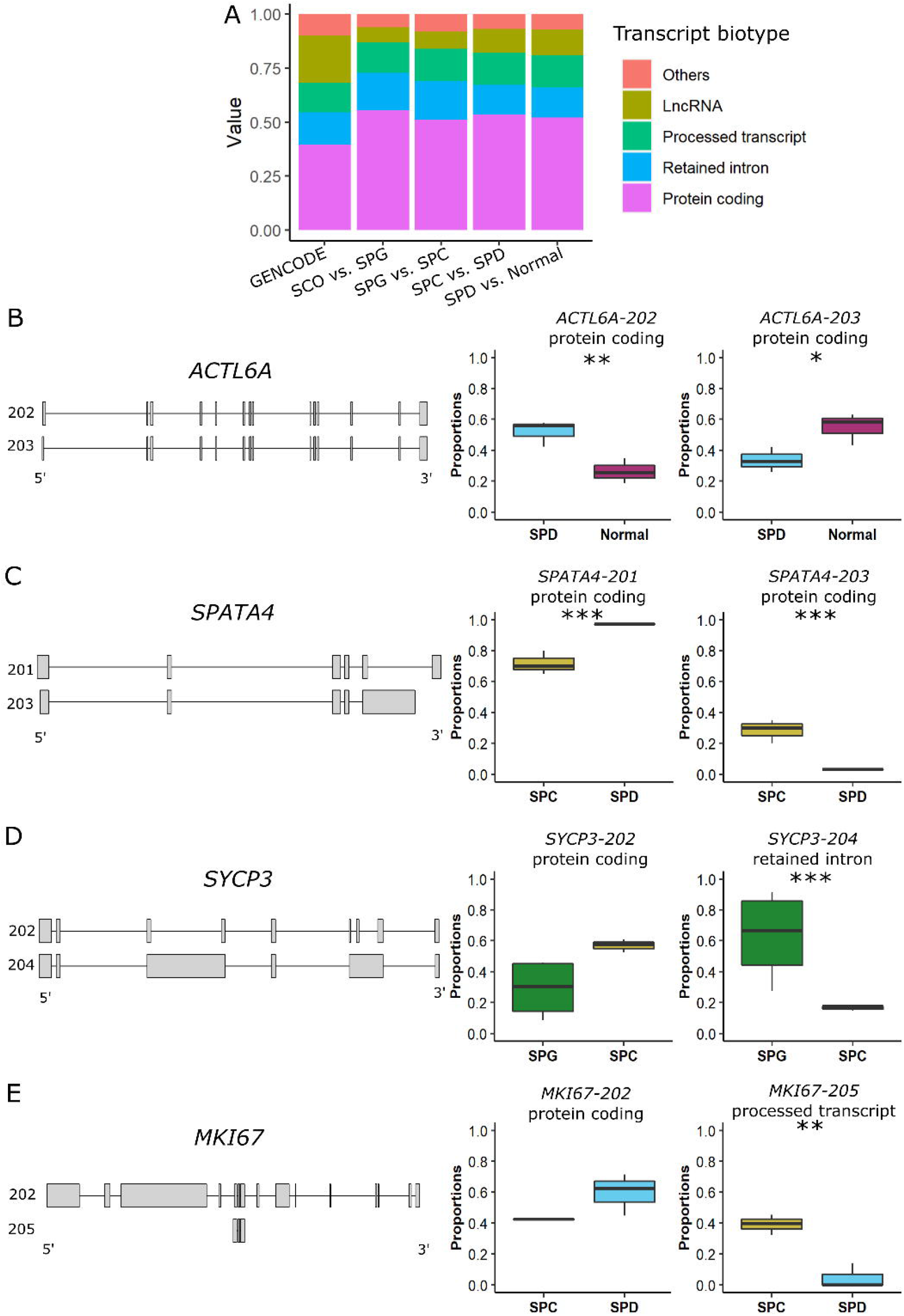
Transcript biotypes with DTU events. (A) Relative amount of different transcript biotypes with DTU events in each of the four group comparisons in comparison to the transcript biotype annotation from the GENCODE release 36 genome annotation based on the GRCh38.p13 genome reference (Frankish *et al*., 2019). (B-E) Schematic illustration of the exons (grey bars) of the transcript isoforms, which predominantly contribute to the relative change in isoform usage (box plots) in (B) *ACTL6A*, (C) *SPATA4*, (D) *SYCP3* and (E) *MKI67. P*-values refer to specific transcripts that significantly drive the change in isoform usage in genes with an overall significant change in transcript usage. * = <0.05, ** = <0.01, *** = <0.001

## Discussion

The study of expression patterns in testis is developing rapidly, however a complete picture of the transcriptome of human germ cells remained unexplored. Here, we demonstrate that the progression of human male germ cell differentiation is accompanied by major transcript dynamics, including germ cell-type dependent transcription and splicing events. The latter resulting in stage-specific transcript isoforms. We found that alternative splicing is mainly uncoupled from the level of gene expression and facilitates a crucial layer of gene regulation in germ cells, especially in the late stages of spermatogenesis.

The differentiation of male germ cells requires cell-specific transcriptional regulation (Guo *et al*., 2018; Hermann *et al*., 2018; Di Persio *et al*., 2021). Previous bulk microarray studies demonstrated that the use of homogeneous human testicular tissues with stage-specific germ cell-arrests allows for the identification of germ cell-specific transcript profiles, thus allowing the unbiased analysis of germ cell populations in their cognate environment (von Kopylow *et al*., 2010; Chalmel *et al*., 2012). Due to the use of microarrays in previous studies, the full spectrum of transcriptome profiles, including isoform information, remained largely unknown.

Our systematic analysis of total RNA from testicular biopsies with well-defined, distinct germ cell compositions revealed significant changes in gene expression (839 to 4,138 DEGs; Supplementary Tables SI, SIII, SIV, SV). Most changes were detected between samples with spermatogonial and spermatocyte arrest, indicating that the entry into meiosis results in a peak of transcriptional activity. Transcripts expressed at this stage are known to be stored for translation at later differentiation stages (Paronetto and Sette, 2010; Wang *et al*., 2020). Among the topmost expressed genes for spermatogonia (2,073), spermatocytes (2,886), round spermatids (2,345) and elongated spermatids (776), we found highly germ cell-specific genes, which to our knowledge were not previously associated with the respective germ cell stages in humans (Supplementary Table SX).

The transcriptional output of a gene depends not only on the level of RNA expression but also on post-transcriptional processing of RNA transcripts, for instance through AS, which allows a single gene to originate different transcripts and potentially different proteins (Baralle and Giudice, 2017). Although it is well known that the testis is an organ with high transcriptome diversity, AS is still understudied in human spermatogenesis. Making use of a powerful bioinformatic technique, the DTU analysis, we were able to study for the first time transcriptome dynamics during human spermatogenesis. Several studies observed discontinuous patterns of transcription throughout murine and human spermatogenesis (Jan *et al*., 2017; Vara *et al*., 2019). In our study, we further characterized the ongoing transcriptional changes during human spermatogenesis by identifying between 1,062 and 2,153 genes whose transcripts were alternatively spliced at different germ cell stages (Supplementary Tables SVI, SVII, SVIII, SIX). Our results indicate that alternative splicing extends the transcriptome diversity in germ cells, which already present high transcriptional activity, as we found that alternative splicing events are more prevalent between the premeiotic and meiotic germ cell stages. As we identified more alternatively spliced genes than changes in gene expression between the spermatid arrest and normal samples, we hypothesize that in the final stage of spermiogenesis transcriptome diversity arises primarily from alternative splicing rather than by changes in gene transcription. In line with this idea are studies in mice showing that genes required for spermiogenesis are already expressed at the beginning of meiosis (da Cruz *et al*., 2016) and that transcription in elongated spermatids is decreased due to the highly compacted chromatin structure (Sassone-Corsi, 2002). Even in the absence of transcriptional activity in the nucleus, stored unprocessed transcripts can maintain translational activity in late stages of germ cell differentiation (Wang *et al*., 2020). Our study demonstrates that alternative splicing is uncoupled from the level of gene expression during human spermatogenesis, as only a minority of genes were both differentially expressed and differentially spliced at each respective germ cell stage. Data on the comparison of DEG and DTU genes in other tissues also revealed that these two gene sets hardly overlap and different molecular functions are enriched (Solovyeva *et al*., 2021). Interestingly, we found that DEGs were enriched for germ cell-specific processes, whereas DTU genes were involved in more general biological processes, suggesting that during human spermatogenesis these functions are predominantly regulated at transcriptional and post-transcriptional level, respectively. We suggest that general processes are uncoupled from the level of gene expression, as these need to be maintained even in transcriptionally silent cells such as later germ cells. By looking more precisely into four DTU genes, we demonstrate the importance of our dataset for further research in the field of male infertility. For example, we were able to reveal that SPD and normal samples express different protein coding transcripts of *ACTL6A*, something that would have been overlooked by conventional DEG analysis. It is also relevant to understand which gene products with potentially different functionality are produced by AS, as it has been shown that this may play a role in the etiology of several diseases (Scotti and Swanson, 2016) such as cancer (Wiesner *et al*., 2015; Vitting-Seerup and Sandelin, 2017). Whether alterations in alternatively spliced transcript expression also plays a role in the pathology of infertility remains to be assessed. We showed that some crucial spermatogenic genes such as *SYCP3* appear to be regulated at both the transcriptional and post-transcriptional levels*. SYCP3* is already expressed as an immature non-coding transcript in SPG samples, whereas the mature transcript is predominantly expressed in SPC samples. A previous study indicated that spermatogonia already express genes required for meiosis (Jan *et al*., 2017), however the mechanism behind this observation was not addressed. In murine spermatogenesis, intron retention ensures timely and stage-depended gene expression (Naro *et al*., 2017). Our data supports the hypothesis that the expression of spermatogenic stage-specific genes might be functionally regulated through alternative splicing by intron retention during human spermatogenesis. Our data strongly highlights the need to further analyze the splicing machinery in human germ cells.

Our whole transcriptome analysis provides an unbiased evaluation of transcriptome dynamics during human spermatogenesis for novel and/or germ cell-specific genes. By not only focusing on protein coding exons but capturing the presence of all alternative transcripts at different stages of human spermatogenesis, our dataset allows to study the role of non-coding pathogenic variants, e.g. in splice sites, by pinpointing the expression and splice isoforms of germ cell-specific transcripts, thereby prospectively improving the genetic diagnosis of male infertility.

## Supporting information

Supplementary Table SII

Supplementary Table SIII

Supplementary Table SIV

Supplementary Table SVI

Supplementary Table SVII

Supplementary Table SVIII

Supplementary Table SVIX

## Author’s roles

Study conception and design: J.G., F.T., N.N. and S.L.; Supervision: N.N., S.L.; Acquisition and evaluation of clinical data: J.F.C., S.K., F.T.; Lab work: N.T., S.D.P., J.W.; Data and bioinformatic analyses: L.M.S.-K., H.K., M.W., T.T., M.D.; Exome analyses/ evaluations: M.J.W., F.T. Writing Original Draft; L.M.S-K., H.K., S.S, N.N., S.L.; All authors were involved in editing, read and approved the final version of the manuscript.

## Acknowledgements

We thank Heidi Kersebom and Elke Kößer for histological evaluation of testicular tissues and we also thank Sabine Forsthoff for excellent support in endocrinological measurements. We thank the service unit Core Facility Genomik of the medical faculty from the University of Muenster for performing the next-generation sequencing. Schematic figure 1A was created with BioRender.com.

## Data availability

The testicular RNA-Seq data of all patients in this study has been deposited in the European Genome-Phenome Archive and is available under EGAS00001006135.

## Supplementary figures and tables

**Figure S1: Levels of gene expression for selected DTU genes.** (Related to figure 6). *P*-values: *** = <0.001

**Table SI: List of DEGs of the SCO vs. SPG group comparison.**

**Table SII: Well-known germ cell markers and related publications.**

**Table SIII: List of DEGs of the SPG vs. SPC group comparison.**

**Table SIV: List of DEGs of the SPC vs. SPD group comparison.**

**Table SV: List of DEGs of the SPD vs. Normal group comparison.**

**Table SVI: List of DTU genes of the SCO vs. SPG group comparison.**

**Table SVII: List of DTU genes of the SPG vs. SPC group comparison.**

**Table SVIII: List of DTU genes of the SPC vs. SPD group comparison.**

**Table SIX: List of DTU genes of the SPD vs. Normal group comparison.**

**Table SX: Novel germ cell marker genes and related publications.**

