## Supplementary figures and images for "Stage-specific gene and transcript dynamics in human male germ cells"

### Figure S1

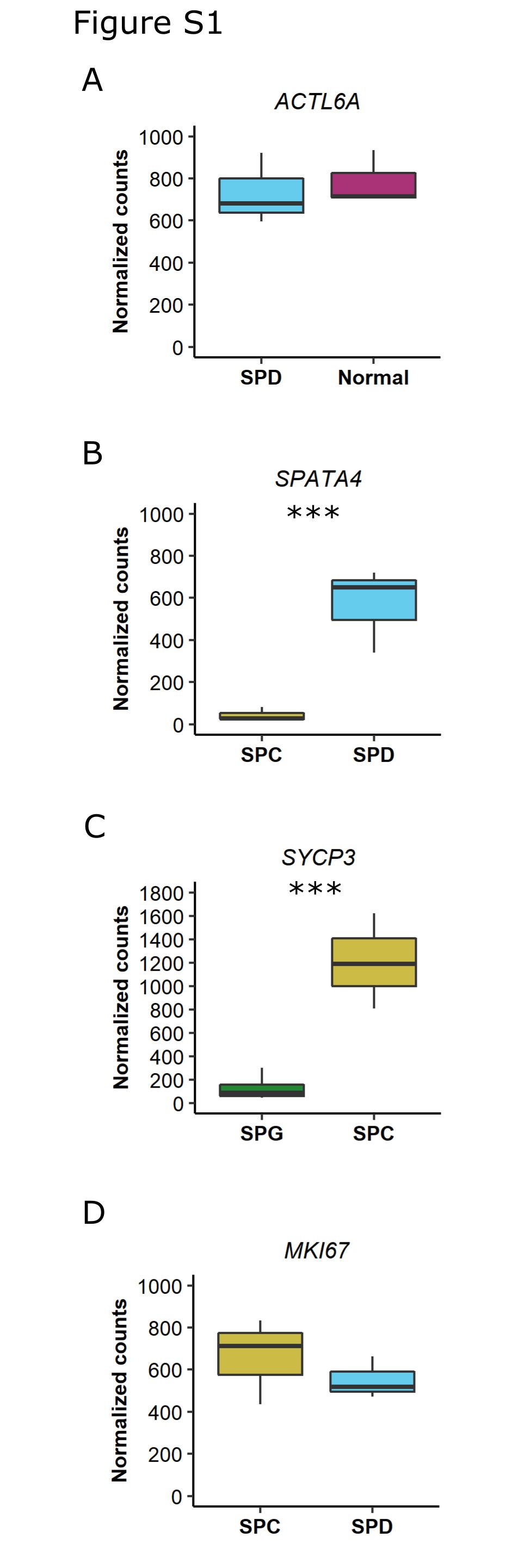
